# Beyond Chemical Similarity: Structure-Agnostic Drug–Drug Interaction Prediction with MeSH Semantics and a Drug–Target–Protein Knowledge Graph

**DOI:** 10.64898/2026.08.10.743843

**Authors:** Ardan Yılmaz, Szymon Szydlik, Golnaz Taheri

## Abstract

**Background:** Adverse drug–drug interactions (DDIs) cause preventable hospitalizations, but exhaustive experimental screening of all drug pairs is infeasible. Many computational predictors rely on SMILES or other molecular representations, limiting their direct applicability to biologics and other non-small-molecule therapeutics. We present a structure-agnostic framework that combines semantic representations derived from Medical Subject Headings (MeSH) with graph-derived topology from a Drug–Target–Protein knowledge graph constructed from DrugBank and UniProt. We further investigate how variation in MeSH annotation depth affects predictive performance.

**Results:** Drugs are grouped according to their deepest MeSH annotation level (Low, Mid, or Deep), and performance is evaluated across the resulting interaction categories in transductive and inductive settings. The Intermediate ontology scope (Low+Mid) provides the most stable performance, while adding Deep-level terms offers limited and inconsistent benefit. Lightweight topological descriptors are integrated with MeSH features through instance-wise, dimension-specific latent-space gating, using curated reliable-negative pairs for supervision. Fusion improves mean performance over the MeSH-only baseline across all six categories in the transductive setting. Under induction, the clearest gains occur for Low–Low interactions (**Δ**AUROC **= 0.056**; **Δ**F1 **= 0.137**) and Low–Mid interactions (**Δ**AUROC **= 0.077**; **Δ**F1 **= 0.114**).

**Conclusions:** MeSH annotation depth is associated with systematic variation in DDI prediction performance that aggregate evaluation can obscure. Graph-derived topology is particularly beneficial when ontology annotations are shallow. The framework provides a common, structure-agnostic representation compatible with both small-molecule and biologic therapeutics and supports first-pass DDI prioritization for subsequent expert assessment.

## 1 Background

**Fig. 1:**
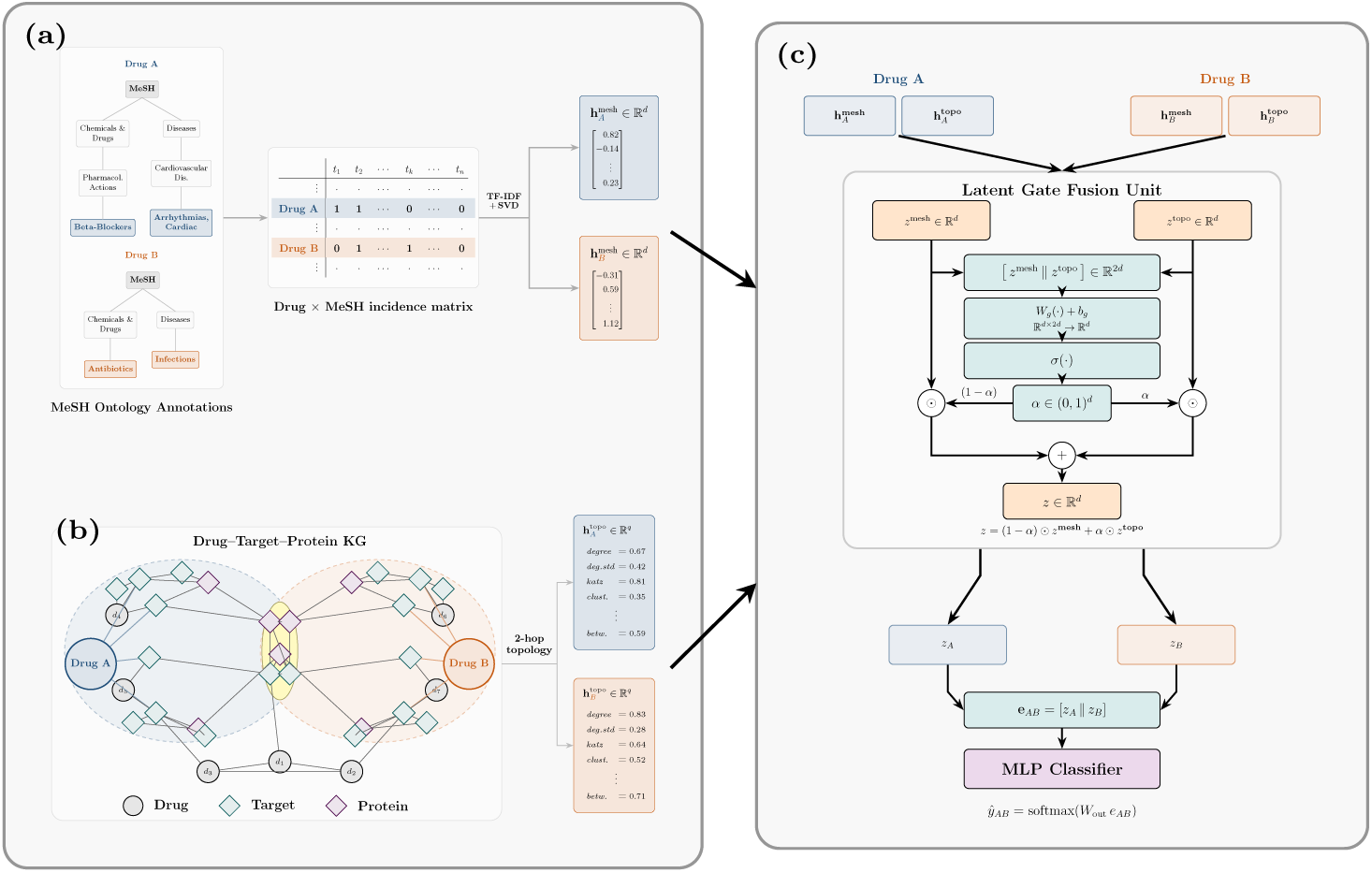
Overview of the proposed multimodal framework for predicting a candidate drug–drug interaction. **(a) MeSH semantics:** each drug is represented using hierarchical MeSH annotations of varying depth. The annotations are converted into a term frequency–inverse document frequency (TF–IDF)-weighted drug–term matrix and reduced using truncated singular value decomposition (SVD). **(b) DTP-KG topology:** lightweight topological descriptors are extracted from each drug’s 2-hop neighborhood in the heterogeneous Drug–Target–Protein knowledge graph. **(c) Fusion and prediction:** the MeSH and topological features are projected into a shared latent space and combined using dimension-wise latent gating. The fused representations of the two drugs are concatenated and passed to an MLP classifier to predict whether the pair interacts.

Polypharmacy—the concurrent use of multiple therapeutics—is now standard practice in the management of chronic and complex diseases. While often clinically necessary, polypharmacy increases the risk of adverse drug–drug interactions (DDIs), some of which cause serious morbidity. The combinatorial space of possible drug pairs is vast, making exhaustive experimental testing logistically infeasible. Moreover, many clinically relevant DDIs are only identified post-market, after drugs are prescribed at scale. Together, these challenges motivate computational DDI prediction as an early safety screen, particularly for new or under-studied therapeutics.

Most machine-learning approaches to DDI prediction formulate the task as binary classification over drug pairs, but two practical limitations remain. First, many pipelines remain chemistry-centric and rely on SMILES-based molecular encodings. This limits their direct applicability to therapeutics that cannot be represented as conventional small molecules, including many biologics and peptide drugs, reducing coverage precisely in settings where prospective screening is most valuable. Although several recent methods explicitly model interactions involving small-molecule and biologic drugs, they generally rely on type-specific molecular representations, such as SMILES or molecular fingerprints for small molecules and amino-acid sequences for biologics [1–3]. Such representations also emphasize molecular structure, while broader biological context, including shared targets, pathways, and protein interaction neighborhoods, may provide complementary information. Second, the task is inherently *positive–unlabeled* (PU): the absence of an interaction annotation does not imply non-interaction. Treating unlabeled pairs as negatives therefore introduces systematic label bias.

To expand DDI prediction beyond small molecules, we analyzed annotation availability across all DrugBank entries (Figure 2), comparing the coverage of candidate modalities between small molecules and biologics. Modalities are ranked by biologic coverage to emphasize availability beyond small molecules. SMILES representations provide near-complete coverage for small molecules in DrugBank, whereas no biologics in the analyzed dataset have a SMILES representation. In contrast, MeSH categories provide broad coverage across both therapeutic classes (59% of small molecules and 53% of biologics), making them a practical shared semantic modality. Interaction-centric annotations (drug–target, drug–drug, enzyme, transporter, carrier, pathway, and reaction relationships) collectively constitute the second highest-ranked modality group in terms of biologic coverage and naturally define a heterogeneous biological network. We therefore focus on these two structured modalities, excluding free-text descriptors to reduce the risk of information leakage and omitting metadata and external cross-references because they fall outside the scope of our semantic and graph-based framework. These observations motivate our choice of two complementary modalities: MeSH and a Drug–Target–Protein Knowledge Graph (DTP-KG).

**Fig. 2:**
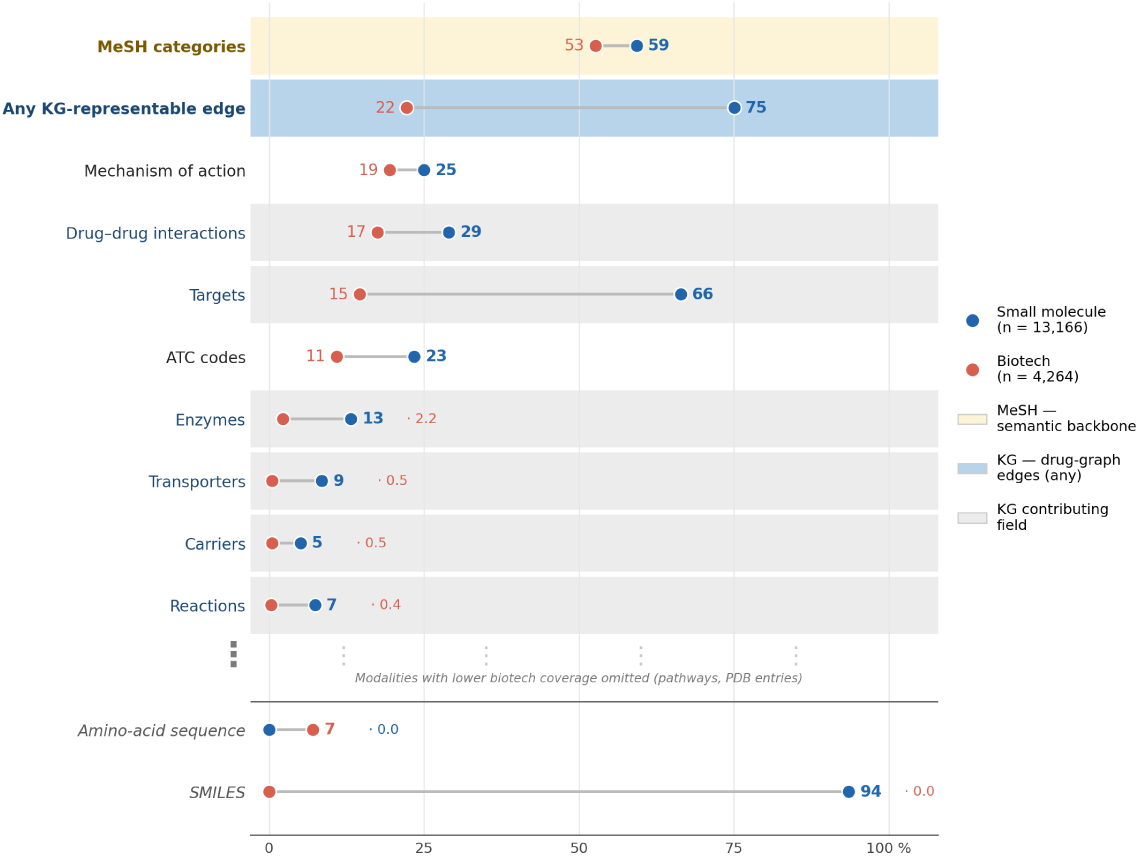
Availability of candidate DrugBank modalities across small molecules and biologics. Rows are ranked by coverage for biologics. Network-representable annotations (e.g., drug–target, drug–drug, enzyme, transporter, carrier, pathway, and reaction relationships) are aggregated into the category Any KG-representable edge. Amino-acid sequence (biologics) and SMILES (small molecules) are shown separately below the divider as modality-specific structural baselines. Free-text descriptions, clinical metadata, and cross-references are excluded.

Building on the earlier DTPPI framework [4], which combined DrugBank drug-category features with topological descriptors from a drug–target–protein–protein interaction network and included both small-molecule and biotechnology drugs, the present study focuses on uneven semantic supervision. We replace broad drug-category features with hierarchical MeSH representations and examine whether graph topology can compensate when ontology annotations are shallow. We further introduce annotation-depth-stratified evaluation, fold-specific graph reconstruction, adaptive latent-space gating, and inductive evaluation on unseen drugs.

Building on these observations, we leverage MeSH, a hierarchical biomedical ontology that annotates therapeutics at varying levels of semantic specificity. However, annotation depth varies substantially across drugs and may partly reflect differences in research and curation intensity: well-characterized drugs often receive deeper annotations, while newer or less extensively studied compounds may have shallower coverage. We refer to this uneven distribution of semantic detail as *hierarchical supervision imbalance*, in which stronger semantic supervision is concentrated on well-characterized drugs. Standard aggregate evaluation under random train/test splits does not make this heterogeneity visible and can obscure weaker performance on shallowly annotated drugs. Such evaluations, therefore, provide a limited view of real-world performance, where predictions are often required for sparsely annotated drugs.

We study this variation systematically by using annotation depth as a proxy for semantic granularity. Specifically, we group drugs based on the deepest MeSH term available (Low, Mid, Deep) and evaluate performance across six interaction categories defined by crossing drug-pair groups. This stratified evaluation makes performance on sparsely annotated drugs explicit, providing a direct assessment of model behavior when semantic metadata are incomplete.

We further incorporate biological network information through a **Drug–Target–Protein Knowledge Graph (DTP-KG)** whose edge types include positive DDIs, drug–target links, and protein–protein interactions. From this graph, we derive lightweight topological features that summarize each drug’s local network context (e.g., connectivity patterns in its 2-hop neighborhood). These graph-derived features can be computed even for drugs with shallow MeSH annotations, providing biological network context that does not rely on deep ontology terms. We note, however, that ontology depth and network connectivity may be correlated because both may be influenced by common research and curation biases: well-characterized drugs often have richer annotations and denser interaction neighborhoods. Nevertheless, our experiments show that incorporating these topological descriptors improves performance, with the largest gains observed for sparsely annotated drugs.

We integrate ontology-derived semantic features and graph-derived topological descriptors by mapping both into a shared latent space and learn instance-wise, dimension-specific fusion weights via latent-space gating. This allows the model to adaptively balance semantic and topological information depending on the available annotations.

Finally, to reduce label bias arising from the *positive–unlabeled* nature of the problem, we employ a curated set of mechanism-supported negative DDI pairs [5]. Rather than treating randomly selected unlabeled pairs as negatives, these curated pairs are used only as supervision labels and are not inserted into the knowledge graph.

Our experiments show that incorporating topological features improves predictive performance, with the clearest gains when one or both drugs lack deep MeSH annotations. These findings indicate that graph-derived biological context provides complementary signal and supports generalization under sparse ontology supervision. As shown in Figure 1, our framework combines MeSH semantics with biological graph topology from DTP-KG and uses latent-space gating to adaptively fuse the two signals, supporting scalable, structure-agnostic first-pass DDI screening even when ontology annotations are sparse. The source code is available at https://github.com/Golnazthr/DTP-KG.

### 1.1 Main Contributions

- We present a structure-agnostic DDI prediction framework that represents small-molecule and biologic therapeutics through a shared combination of MeSH semantics and Drug–Target–Protein Knowledge Graph topology, without requiring SMILES or amino-acid-sequence encodings.
- We identify and systematically analyze performance variation associated with heterogeneous MeSH annotation depth, which we term *hierarchical supervision imbalance*, and provide a stratified evaluation framework for studying performance under varying levels of semantic supervision.
- We use lightweight, leakage-aware topological descriptors derived from DTP-KG and integrate them with ontology-derived semantics through instance-wise, dimension-specific latent-space gating.
- We reduce label bias by training with curated reliable negative interactions and evaluate performance across knowledge-depth strata and inductive splits to expose model behavior under sparse-annotation and unseen-drug conditions.

### 1.2 Related Work

Early DDI prediction methods relied on chemical similarity and molecular descriptors. More recently, graph neural networks and transformer-based encoders over molecular graphs have become common for (typed) interaction prediction [6–8]. While these models can be highly effective, many remain chemistry-centric: their applicability depends on the availability of conventional molecular representations, and mechanistic biological context is frequently underrepresented.

A smaller body of work has explicitly considered interactions involving biologic or biotechnology drugs. Multi-SBI combines SMILES-derived molecular fingerprints and sequence encoders with heterogeneous network-topology features, and applies positive–unlabeled sampling to identify reliable negative examples [1]. CB-TIP jointly learns from molecular graphs and an interaction graph containing drug–drug, drug–protein, and protein–protein relations to predict interaction events involving chemical and biotechnology drugs [2]. More recently, BSI-Net combined SMILES-derived molecular graphs for small molecules with protein-language-model embeddings for biotechnology drugs [3]. These methods demonstrate the feasibility of predicting interactions involving biologic therapeutics, but they rely on modality-specific molecular inputs, such as SMILES, molecular fingerprints, or amino-acid sequences, rather than a common structure-independent representation across therapeutic classes.

To incorporate biological and relational information, knowledge-graph-based methods model drug, target, protein, disease, and interaction relations using neighborhood aggregation, subgraph reasoning, contrastive learning, or heterogeneous GNNs. KGNN showed that drug representations can be learned from biomedical knowledge-graph neighborhoods without requiring chemical structures [9]. More recent approaches include KnowDDI, which learns pair-specific and interpretable knowledge subgraphs from a biomedical knowledge graph [10], and KG-CLDDI, which combines knowledge-graph aggregation with topological information from the DDI graph through contrastive learning [11]. Although these approaches highlight the value of relational structure, they do not explicitly stratify performance according to ontology annotation depth or examine whether graph context compensates for uneven semantic supervision across drugs.

The most direct precursor to the present work is the DTPPI framework [4], which combines DrugBank category features with topological descriptors derived from a weighted drug–target–protein–protein interaction network and includes both small-molecule and biotechnology drugs. Compared with DTPPI, the present study replaces broad drug-category features with ancestor-closed hierarchical MeSH representations, introduces annotation-depth-stratified evaluation and fold-specific graph reconstruction, and uses instance-wise latent-space gating to integrate semantic and topological information. It further evaluates generalization to drugs that are held out entirely during training.

A separate but related issue is label quality. Many DDI models treat unlabeled drug pairs as negatives, implicitly equating missing annotations with non-interaction and thereby introducing systematic label bias. Positive–unlabeled objectives and sampling strategies have been proposed to mitigate this issue [12**?**], while curated mechanism-supported negative interactions provide an alternative source of supervision [5].

Table 1 summarizes representative methods most closely related to the present study in terms of therapeutic scope, input representation, and evaluation focus.

**Table 1:**
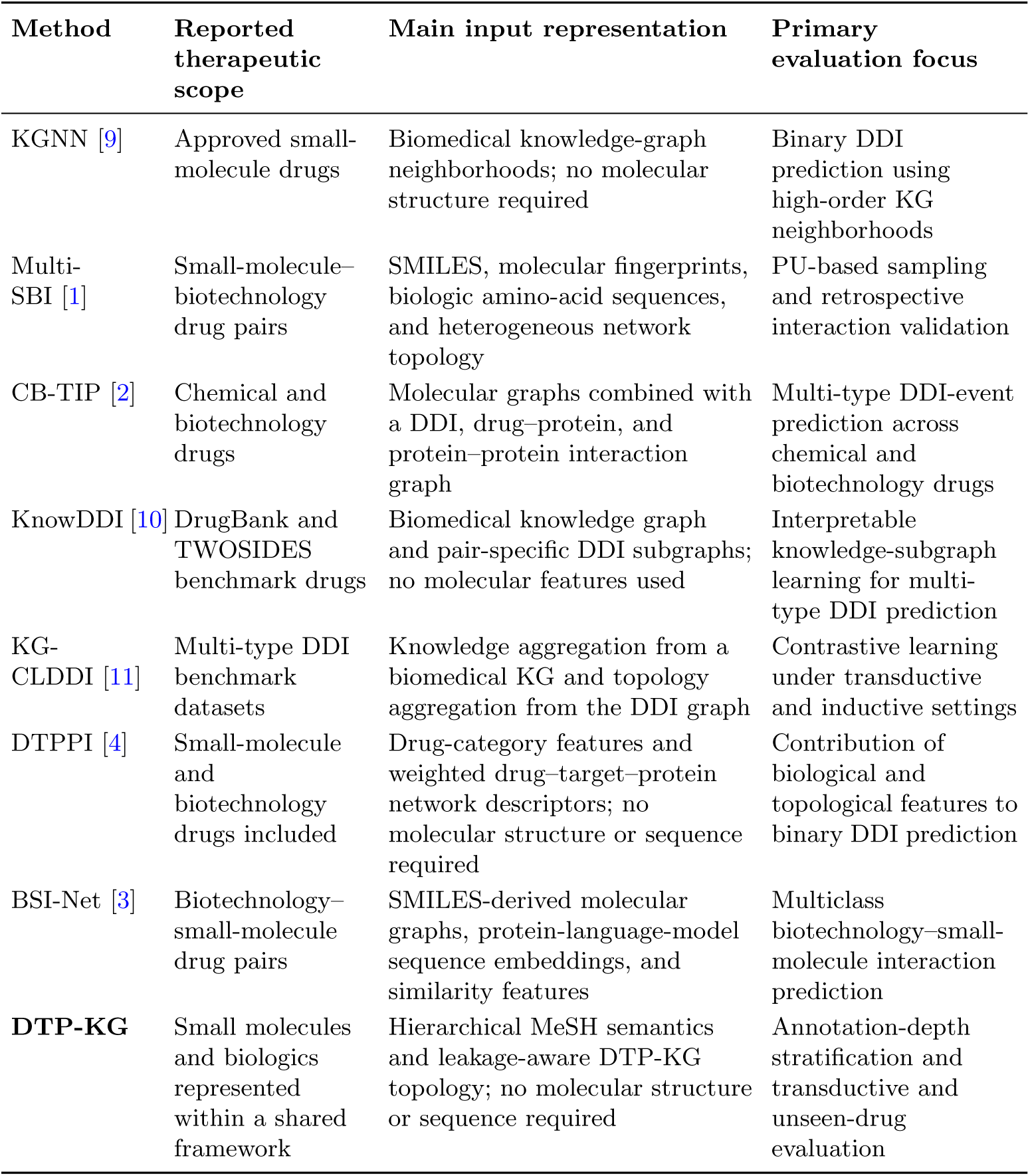
Qualitative comparison of representative DDI prediction approaches related to the present study. Numerical performance values are not compared because the methods use different datasets, interaction definitions, negative-sampling procedures, and evaluation protocols.

Relative to these lines of work, our framework focuses on (i) representation coverage beyond small molecules through a shared structure-agnostic feature space, (ii) performance variation associated with heterogeneous MeSH annotation depth, and topology and adaptive latent-space fusion, and evaluates the resulting model across annotation-depth strata and inductive unseen-drug splits.

## 2 Methods

Figure 1 summarizes the proposed DDI screening pipeline. For a candidate drug pair (*d_i_, d_j_*), we extract (i) semantic descriptors from MeSH and (ii) lightweight topological descriptors from a Drug–Target–Protein knowledge graph (DTP-KG). We project both modalities into a shared latent space and fuse them via latent-space gating, after which an MLP predicts whether the pair corresponds to a known DDI.

### 2.1 Problem Definition and Data Sources

Given a drug pair (*d_i_, d_j_*), we predict a binary label *y_ij_*∈ {0, 1}, where *y_ij_* = 1 denotes a documented positive DDI and *y_ij_* = 0 denotes a pair reported as a reliable negative by Zheng et al. [5]. We construct our dataset from DrugBank [13], Zheng et al. [5], and UniProt [14]. Positive DDIs, drug–target interactions, and MeSH descriptors [15] were obtained from DrugBank 5.1.13. Protein–protein interactions were obtained from UniProt release 2025 03, and reliable negative DDI pairs were taken from the supplementary data of Zheng et al. [5].

#### Drug selection and preprocessing

The candidate drug set is defined by the union of documented positive DDIs from DrugBank and the reliable negative interactions reported by Zheng et al. [5]. Drug-Bank primary accessions are used as the common identifier across all resources. MeSH annotations are obtained directly from DrugBank and expanded with all ancestor terms in the MeSH hierarchy, while drug–target and protein–protein interactions are integrated through their shared UniProt identifiers. Drugs lacking valid MeSH annotations are excluded. Within each ontology scope, MeSH terms annotating fewer than two drugs or more than a predefined threshold are removed, thereby discarding both overly specific and overly generic terms (Figure 3). Drugs left without any remaining annotations after filtering are excluded, and only drug pairs for which both drugs satisfy these criteria are retained. No additional filtering based on graph connectivity is performed beyond requiring each retained drug to be represented in the constructed drug–target–protein network.

**Fig. 3:**
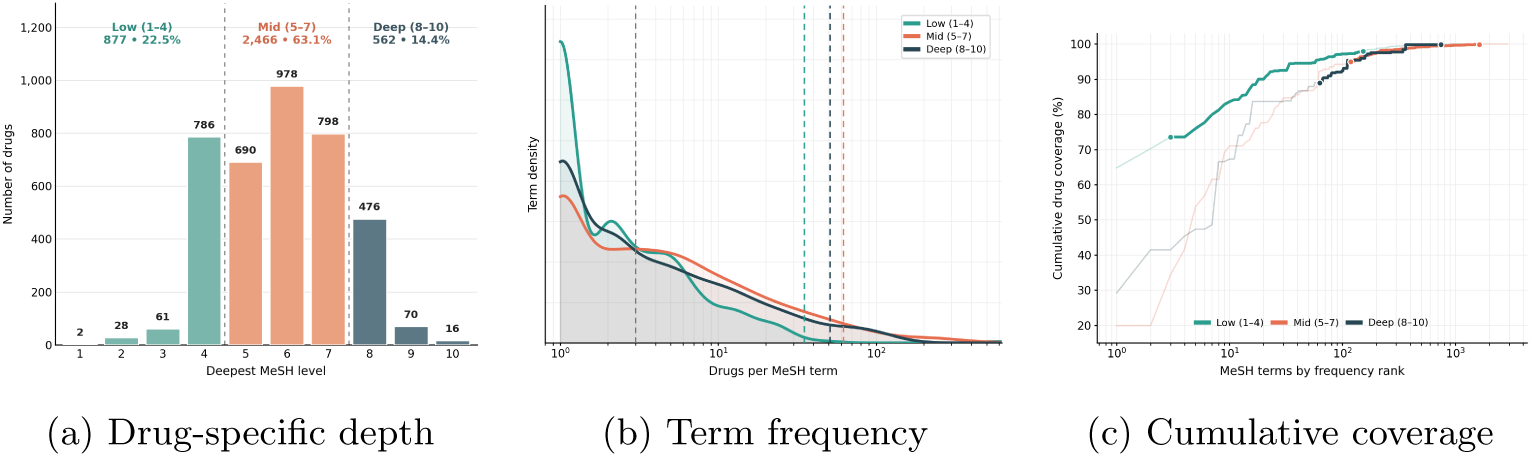
MeSH annotation depth and term-selection strategy. **(a)** Distribution of drugs according to their deepest assigned MeSH hierarchy level. **(b)** MeSH term-frequency distributions for the Low (teal), Mid (orange), and Deep (dark blue) knowledge bands. The grey dashed line indicates the shared lower-frequency threshold of three drugs per term, while the colored dashed lines indicate the corresponding upper-frequency thresholds for the Low, Mid, and Deep bands. **(c)** Cumulative drug coverage as terms are added in decreasing order of frequency. Colors correspond to the same knowledge bands as in **(b)**.

#### Final dataset

Following preprocessing, the positive interaction set is randomly downsampled to match the number of reliable negatives, yielding balanced datasets for each ontology scope. The intermediate ontology, used in the main experiments, contains 1,667 drugs and 10,782 drug pairs (5,391 positive and 5,391 reliable negative). The corresponding drug–target–protein network comprises 17,976 nodes (3,263 drugs, 2,886 targets, and 11,827 proteins) and 268,307 edges.

#### Node and edge types

We build an undirected heterogeneous graph G = (*V, E*) with three node types: drugs (*D*), targets (*T*), and other proteins (*P*), such that *V* = *D*∪*T* ∪*P*. Targets are proteins that appear in DrugBank’s drug–target interaction tables; proteins included through UniProt interactions that do not appear as DrugBank targets are assigned to *P*. The edge set comprises five typed relations:

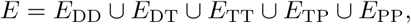

where *E*_DD_ contains positive DDIs, *E*_DT_ contains drug–target links, and *E*_TT_, *E*_TP_, and *E*_PP_ are derived from protein–protein interactions between targets and/or other proteins. Reliable negative DDIs are used as supervision for classification but are not inserted into the graph as edges.

The resulting graph contains 3,263 drug nodes, 2,886 target nodes, and 11,827 additional protein nodes. The labeled pair set contains 720,172 positive DDIs and 5,517 reliable negative pairs.

### 2.2 MeSH Feature Engineering

MeSH is a hierarchical controlled vocabulary curated by the U.S. National Library of Medicine. DrugBank provides MeSH terms from the *D* branch, which represents chemical and pharmacological concepts. Greater tree depth generally corresponds to more specific concepts [16].

#### Hierarchy closure

DrugBank typically reports the most specific MeSH terms assigned to each drug. For each drug *d*, we therefore construct the ancestor-closed set *M* (*d*) by propagating its assigned terms upward through the MeSH hierarchy. This preserves valid higher-level semantics when supervision is restricted to shallower ontology levels, rather than discarding informative ancestor terms.

#### Knowledge-level variation and ontology-scope variants

MeSH depth varies substantially across drugs. For each drug, we record the deepest assigned MeSH hierarchy level and stratify drugs into **Low** (levels 1–4), **Mid** (levels 5–7), and **Deep** (levels 8–10), as shown in Figure 3(a). This stratification induces six interaction categories: Low–Low, Low–Mid, Low–Deep, Mid–Mid, Mid–Deep, and Deep–Deep.

We further define three ontology-scope variants with increasing semantic depth: **Shallow**, containing Low-level terms only; **Intermediate**, containing Low- and Mid-level terms; and **Full**, containing terms from all three levels. These variants control the depth of MeSH information available to the model while retaining the same drug-pair evaluation structure.

#### Frequency filtering and compact embeddings

To limit noise from extremely rare or overly generic terms, we filter MeSH terms according to their annotation frequencies, guided by the frequency and cumulative-coverage profiles shown in Figures 3(b) and 3(c). We apply a shared lower-frequency threshold of three drugs per term together with band-specific upper-frequency thresholds. We then construct a binary drug–term incidence matrix over *M* (*d*), apply term frequency–inverse document frequency (TF–IDF) weighting, and obtain compact embeddings through truncated singular value decomposition (SVD). This procedure results in vocabularies of 1,416, 2,128, and 2,183 MeSH terms for the Shallow, Intermediate, and Full variants, respectively. Each representation is reduced to 128 dimensions using truncated SVD.

### 2.3 Handling Class Imbalance

The reliable-negative set is substantially smaller than the positive set (720,172 positive pairs versus 5,517 reliable negative pairs). To avoid treating additional unlabeled drug pairs as negative examples, we downsample the positive class within each training fold.

Data augmentation is widely used in vision and language, where operations such as image rotation or controlled noise injection can preserve the original label. There is no directly analogous, biologically grounded transformation for drug pairs. Small perturbations in feature or embedding space lack a clear pharmacological interpretation and may produce implausible examples. We therefore do not use synthetic data augmentation. We did not systematically evaluate alternative imbalance-mitigation strategies, such as class-weighted or cost-sensitive loss functions; this is considered a limitation of the present study.

### 2.4 Topological Features from the DTP-KG

We complement MeSH representations with topological descriptors derived from the DTP-KG, which encodes known DDIs, drug–target interactions, and protein–protein interactions.

Computing topological features once on the complete graph and reusing them across folds could introduce information leakage because the resulting descriptors may encode properties of held-out DDI edges. To reduce this risk, DDI edges assigned to the test fold are removed before feature extraction, and all graph-derived features are recomputed independently within each cross-validation fold.

While larger ego networks may capture more extensive biological and topological context, they come at considerable computational cost because the number of reachable nodes and edges increases rapidly with neighborhood size. The extraction of higher-order descriptors therefore becomes increasingly expensive. For example, local clustering can require *O*(*d*(*v*)^2^) operations, while iterative Katz centrality requires approximately *O*(*T* |*E*_ego_|) operations, where *T* is the number of power iterations and |*E*_ego_| is the number of edges in the ego network. We therefore restrict feature extraction to each drug’s 2-hop ego network, providing a practical compromise between information content, computational efficiency, and interpretability.

#### Per-drug descriptors

Within each drug’s 2-hop ego subgraph, we compute lightweight topological descriptors that quantify its local connectivity and position relative to targets, neighboring proteins, and other drugs. The following descriptors are used:

- **Degree-based connectivity:** total degree, drug-neighbor degree, and protein-neighbor degree, capturing the density and composition of the drug’s immediate connections.
- **Neighbor-degree statistics:** minimum, maximum, mean, and standard deviation of neighbor degrees, describing the connectivity profile and heterogeneity of the local neighborhood.
- **Clustering coefficient:** local triangle density, providing a measure of neighborhood cohesion.
- **Boundary ratio:** the fraction of edges that leave the ego subgraph, indicating whether the drug lies within a locally cohesive region or near the interface between network regions.
- **Closeness centrality:** the inverse of the average shortest-path distance from the drug to reachable nodes in its ego subgraph, measuring its local accessibility.
- **Local Katz centrality:** influence through attenuated walks within the ego subgraph, capturing connectivity beyond immediate neighbors.
- **Local betweenness centrality:** the extent to which the drug lies on shortest paths between other nodes in its ego subgraph, capturing its local bridging role.

### 2.5 Multimodal Fusion via Latent-Space Gating

MeSH and topology features differ in scale and geometry; näıve concatenation may allow the higher-dimensional MeSH representation to dominate. We therefore encode both modalities into a shared *d*-dimensional latent space, where *d* = 128, and learn an instance-specific gate that balances their contributions for each drug.

For a drug *u*, let 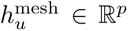 and 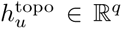 denote its MeSH and topological features, respectively, where *p* = 128 and *q* = 12. We encode each modality into the shared latent space:

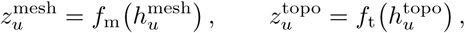

where *f*_m_ and *f*_t_ are MLP encoders mapping their respective inputs into ℝ*^d^*, yielding 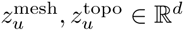. Both encoders share the same architecture: a two-layer MLP with a 256-dimensional hidden layer, ReLU activation, dropout (0.1), and a linear projection to the shared latent space, followed by layer normalization. Thus, *f*_m_ maps ℝ^128^ → ℝ^256^ →ℝ^128^, whereas *f*_t_ maps ℝ^12^ →ℝ^256^ →ℝ^128^.

We then compute a dimension-wise gating vector:

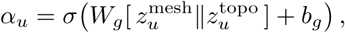

where *W_g_* ∈ ℝ*^d^*^×2*d*^, *b_g_* ∈ ℝ*^d^*, and *α_u_* ∈ (0, 1)*^d^*. The gate consists of a single linear layer followed by a sigmoid, with no hidden layer or dropout.

The two modalities are fused element-wise:

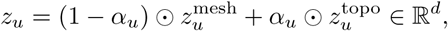

where ⊙ denotes element-wise multiplication.

To score a candidate interaction between drugs *u* and *v*, we concatenate their fused representations and pass them to a three-layer MLP classifier:

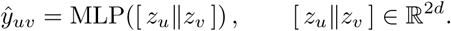

The prediction head has architecture 256 → 256 → 128 → 1, with ReLU activations and dropout (0.1) after the first two linear layers. The network outputs a raw logit, while the sigmoid activation is incorporated into the binary cross-entropy objective (`BCEWithLogitsLoss`) for improved numerical stability.

The *MeSH-only* baseline omits the modality encoders and the latent gate. It passes the raw MeSH pair representation 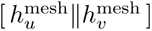 directly to a prediction head that is architecturally identical to that of the fusion model. Since both pair representations have dimensionality 256, the two models share the same three-layer prediction head (256 → 256 → 128 → 1), the same ReLU activations, the same dropout rate (0.1), and the same raw-logit output. Consequently, any performance difference between the two models is attributable solely to the learned multimodal representation rather than to differences in prediction-head capacity.

### 2.6 Experimental Setup

Unless otherwise stated, experiments use 5-fold cross-validation. Analyses that use a different number of repetitions report this explicitly in the relevant subsection.

Models are trained using the Adam optimizer with a learning rate of 10^−3^ and weight decay of 10^−5^, minimizing binary cross-entropy on the raw logits (`BCEWithLogitsLoss`). Dropout with rate 0.1 is applied after each hidden layer, and layer normalization is used at the encoder outputs. Training is performed for up to 100 epochs (50 epochs for the inductive MeSH-only baseline), with early stopping based on the validation loss using a patience of 5 epochs. The checkpoint achieving the lowest validation loss is retained for evaluation.

We evaluate performance using Accuracy, Precision, Recall, F1 score, and area under the receiver operating characteristic curve (AUROC). Results are reported either as absolute values or as differences between paired training regimes, depending on the experiment. Statistical significance is assessed using paired tests on matched fold-and-run results, with the test used for each analysis specified in the corresponding subsection.

Evaluation proceeds in two stages followed by an inductive test. First, we analyze the effect of semantic depth by comparing the Shallow, Intermediate, and Full ontology-scope variants across the six MeSH-depth interaction categories. Under the selected ontology scope, we then assess whether adding topological features through latent-space gating produces consistent gains over the MeSH-only baseline.

Finally, we assess inductive generalization by holding out 103 drugs entirely from model training. Held-out drugs are sampled with stratification by MeSH knowledge category and graph degree. All labeled DDI pairs involving at least one held-out drug are reserved for the inductive test set. A stratified 15% validation split of the training pairs is used exclusively for early stopping and checkpoint selection. To prevent interaction-label leakage, normalization parameters are estimated using the training drugs only. Non-DDI biological relations required to construct representations for the held-out drugs are retained.

## 3 Results

### 3.1 Effect of MeSH Ontology Scope

We investigate how the depth of ontology information available during training affects predictive performance across the six MeSH-depth interaction categories. Specifically, we compare three MeSH ontology scopes: *Shallow* (Low-level terms only), *Intermediate* (Low+Mid levels), and *Full* (Low+Mid+Deep levels). These ontology-scope variants control the amount of semantic information available to the model while keeping the evaluation splits fixed.

Figure 4 summarizes the AUROC and F1 differences between ontology scopes across the six interaction categories.

**Fig. 4:**
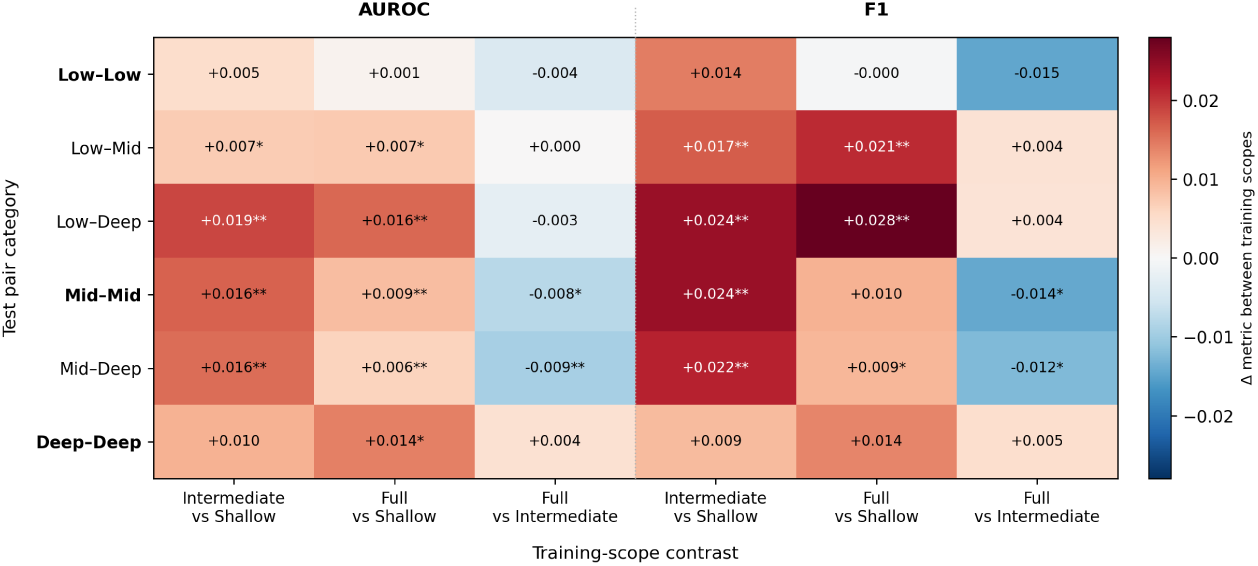
Performance differences between MeSH ontology scopes. Rows correspond to interaction categories defined by the MeSH-depth combination of the interacting drugs, while columns compare *Intermediate vs. Shallow*, *Full vs. Shallow*, and *Full vs. Intermediate*. Positive values indicate that the first-listed ontology scope outperforms the second. Asterisks denote paired *t*-tests on fold- and seed-matched results across 3 seeds × 5 folds (^∗^*p <* 0.05, ^∗∗^*p <* 0.01).

#### Key observations

The most consistent gains arise when expanding the ontology scope from *Shallow* to *Intermediate*. As shown by the *Intermediate vs. Shallow* comparison, AUROC and F1 improve across nearly all interaction categories, indicating that Mid-level MeSH terms provide substantial additional semantic information beyond shallow annotations.

In contrast, the *Full vs. Intermediate* comparison shows only marginal improvements and occasional performance decreases. These results indicate that most of the useful semantic signal is already captured by the *Intermediate* ontology scope, while including Deep-level terms provides limited and inconsistent additional benefit.

We therefore use the *Intermediate* ontology scope as the **default semantic feature set** throughout the remainder of the study.

### 3.2 Fusion vs. MeSH-Only Baseline

Next, we quantify the contribution of topology by comparing the MeSH-only baseline with the fusion model, which combines MeSH and topological features through latent-space gating. We first evaluate the models in the *transductive setting*, where interaction pairs are divided between training and test sets and individual drugs may appear in both sets.

Both models are trained and evaluated on identical folds, allowing matched comparisons. The transductive results are shown in the left column of Figure 5: AUROC in panel (a), F1 in panel (c), and the per-metric improvement (Δ = Fusion−MeSH-only) in panel (e).

**Fig. 5:**
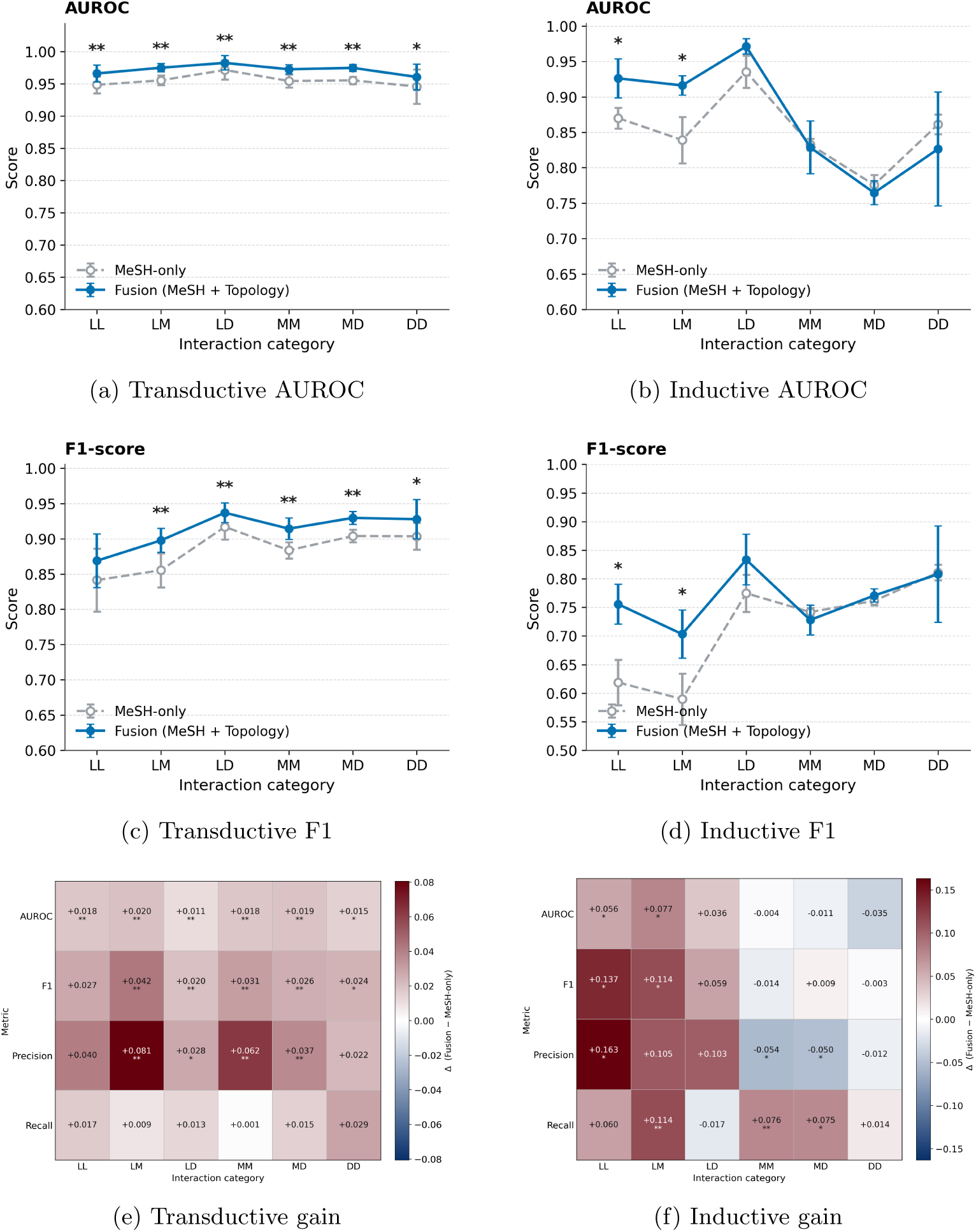
Performance of the MeSH-only baseline and multimodal fusion (MeSH + topology) in transductive and inductive settings. The left column shows pair-level transductive evaluation, while the right column shows drug-level inductive evaluation on held-out drugs. The top and middle rows report AUROC and F1; the bottom row shows the mean difference (Δ = Fusion − MeSH-only). LL, LM, LD, MM, MD, and DD denote the six MeSH-depth pair categories. Points show means, error bars indicate one standard deviation across folds and seeds, and asterisks indicate paired *t*-tests (^∗^*p <* 0.05, ^∗∗^*p <* 0.01).

#### Key observations

1. Despite the strong performance of the MeSH-only baseline, with a mean AUROC of 0.955, incorporating topology improves mean predictive performance across all six interaction categories. Fusion improves AUROC for Low–Low (LL), Low–Mid (LM), Low–Deep (LD), Mid–Mid (MM), Mid–Deep (MD), and Deep–Deep (DD) interactions. The transductive gain heatmap also shows positive mean differences across all reported metric–category combinations. Statistically significant improvements are observed for the majority of the evaluation metrics, particularly for LM, MM, and MD interactions. Improvements for LL and DD are generally smaller or less consistently significant. These results indicate that topological information provides complementary predictive signal beyond the MeSH-derived semantic representation.
2. Recall is an exception to the overall pattern. Its mean improvement is smaller (Δ = +0.014), and the differences do not reach statistical significance in any interaction category. The larger gains in AUROC, Precision, and F1 indicate that the transductive benefit of topology is driven more by improved discrimination and precision than by a substantial increase in sensitivity.

### 3.3 Inductive Generalization to Unseen Drugs

To assess generalization to unseen drugs, we construct an inductive split by holding out 103 drugs from all training pairs. In the transductive setting, the data are partitioned at the interaction-pair level, so the same drug may appear in both training and test sets through different pairs. The inductive setting instead partitions the data at the drug level: none of the held-out drugs appears in a training pair, and every test pair contains at least one held-out drug.

Held-out drugs are sampled with stratification by MeSH knowledge category (Low, Mid, Deep) and KG degree category (low, mid, high) to approximately preserve the corresponding distributions. All labeled DDI pairs involving at least one held-out drug are reserved for the inductive test set.

To reduce interaction-label leakage, positive DDI edges incident to the held-out drugs are removed before topological feature computation, and normalization statistics are estimated using the training drugs only. The inductive results are shown in the right column of Figure 5: AUROC in panel (b), F1 in panel (d), and the per-metric gain in panel (f).

#### Key observations

1. Overall performance is lower and more variable than in the transductive setting. AUROC ranges from approximately 0.77 to 0.97 across interaction categories, compared with values concentrated around 0.95 in the transductive evaluation, and the inductive results exhibit substantially wider error bars. This pattern is consistent with the greater difficulty of predicting interactions involving unseen drugs.
2. The clearest benefit of fusion is observed in categories involving at least one shallowly annotated (*Low*) drug. Fusion significantly improves both AUROC and F1 for Low–Low interactions (ΔAUROC = +0.056, ΔF1 = +0.137; *p <* 0.05) and Low–Mid interactions (ΔAUROC = +0.077, ΔF1 = +0.114; *p <* 0.05). These are the categories with the least detailed MeSH supervision, indicating that topology provides complementary information when ontology annotations are shallow.
3. When neither drug is shallowly annotated, the net gains in AUROC and F1 are generally smaller. For Mid–Mid, Mid–Deep, and Deep–Deep interactions, the results do not show the same consistent improvement observed for Low–Low and Low–Mid pairs. This suggests that the MeSH-derived representation is already highly informative in these categories, leaving less room for additional improvement from topology.
4. Deep–Deep is the only interaction category with a negative mean AUROC difference under fusion (ΔAUROC = −0.035). The difference is not statistically significant and should therefore not be interpreted as evidence of a systematic performance decrease.
5. In contrast to the transductive setting, fusion significantly increases Recall under induction for Low–Mid, Mid–Mid, and Mid–Deep interactions (*p <* 0.05). For Mid–Mid and Mid–Deep interactions, however, the increase in Recall is accompanied by a significant reduction in Precision, while AUROC and F1 remain effectively unchanged. This pattern represents a precision–recall trade-off rather than a consistent overall improvement.

## 4 Ablation

### 4.1 Ablation I: Evaluating Fusion Strategies

We compare latent-space gating with simpler fusion schemes to assess the effectiveness of different strategies for combining MeSH-derived semantics and topological descriptors. All variants are evaluated using identical data splits (stratified 5-fold cross-validation with five random seeds) and the same downstream MLP prediction head. Direct concatenation combines the two feature vectors in the input space, while the weighted variants multiply the topological feature vector by either a fixed or a learned global scalar *α* before concatenation. For each fusion strategy, we perform a modest hyperparameter search and report the best-performing configuration. All variants use the same fold-specific leakage-prevention procedure described previously, including the removal of test DDI edges and recomputation of graph-derived features.

Table 2 summarizes the performance of the evaluated fusion schemes. Direct concatenation and weighted concatenation, using either a fixed or a learned global weight, yield relatively small improvements over the MeSH-only baseline (Δ = 0.0061–0.0094). None of these comparisons with the MeSH-only baseline reaches the nominal significance threshold according to DeLong’s test.

**Table 2:**
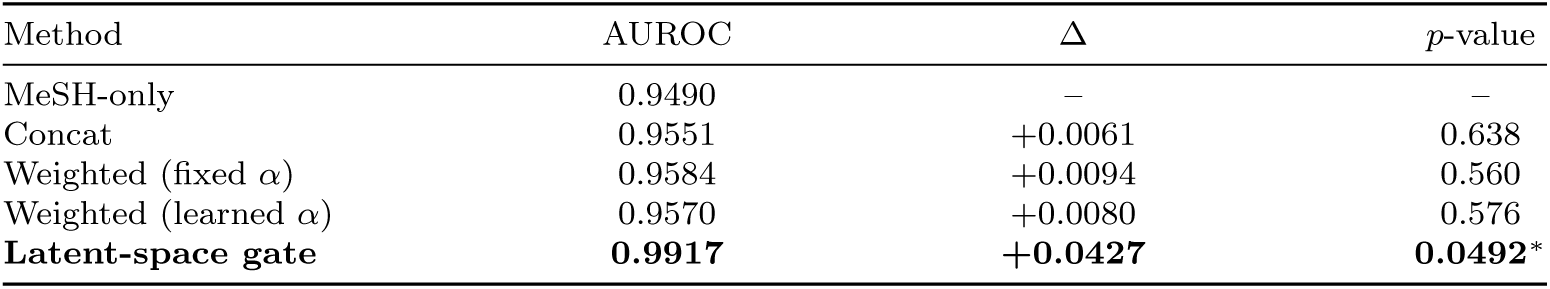
Comparison of fusion strategies. AUROC values are calculated from pooled out-of-fold predictions obtained using five random seeds and five stratified cross-validation folds per seed, with identical folds across methods. The *p*-values are obtained using DeLong’s test and compare each fusion strategy with the MeSH-only baseline. Δ denotes the AUROC difference relative to the MeSH-only baseline; ^∗^ denotes nominal *p <* 0.05.

The proposed latent-space gating strategy achieves the highest AUROC among the evaluated fusion schemes, improving AUROC from 0.9490 to 0.9917 (Δ = 0.0427). Its comparison with the MeSH-only baseline is the only one to reach the nominal threshold of *p <* 0.05 (*p* = 0.0492). Under this experimental setup, the simpler input-space fusion strategies do not produce a comparable improvement. These results suggest that instance-wise, dimension-specific fusion is more effective than direct concatenation or global feature weighting for integrating the two modalities.

The pooled AUROC values in Table 2 are calculated differently from the category-wise mean AUROC values reported in Figure 5. The two sets of values should therefore not be compared directly.

### 4.2 Ablation II: Contribution of Topological Feature Groups

We next assess the contribution of different groups of topological descriptors. To keep the analysis interpretable, we ablate coherent feature groups rather than individual descriptors, for which the effects can be difficult to attribute because of correlations among related features. All experiments in this subsection are performed in a topology-only setting, without MeSH supervision, to evaluate the network-topological signal in isolation and avoid compensatory effects from the semantic modality.

We define the following groups of related topological features:

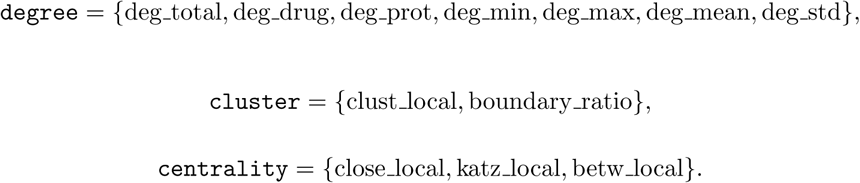

We train and evaluate four *topology-only configurations*: (i) a model using all topological features, and three grouped ablations in which (ii) `degree`, (iii) `cluster`, or (iv) centrality features are removed. Table 3 reports AUROC and F1, together with the corresponding changes relative to the full topology model.

**Table 3:**
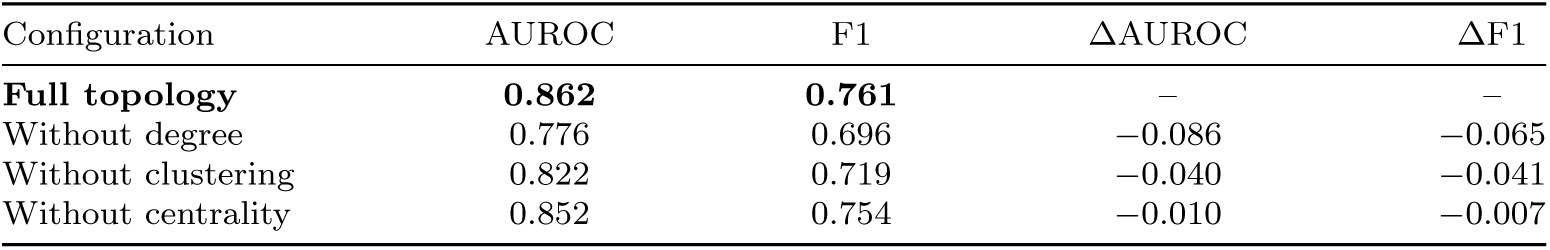
Topology-only feature-group ablation. Models are trained using topological descriptors alone, without MeSH supervision. Reported AUROC and F1 values are averaged over five cross-validation folds and six interaction categories. Δ denotes the change relative to the full topology model.

Removing the degree feature group produces the largest observed decrease in both AUROC and F1, indicating that degree-related descriptors provide the largest contribution to the topology-only model in this experiment. Removing the cluster or centrality groups results in smaller performance decreases, suggesting that these descriptors provide additional but comparatively smaller contributions.

## 5 Case Study: Interpreting Topology-Only Predictions

To provide qualitative intuition about the information captured by the topological descriptors, we train a topology-only model and visualize the 2-hop ego neighborhoods of two correctly classified test pairs: a documented positive DDI assigned a high interaction score and a curated reliable-negative pair assigned a low interaction score. The positive example is mirabegron (DrugBank: DB08893) and antrafenine (DrugBank: DB01419), which has reference label *y* = 1 and receives a predicted interaction score of 1.00. The negative example is ethosuximide (DrugBank: DB00593) and *γ*-aminobutyric acid (DrugBank: DB02530), which has reference label *y* = 0 and receives a predicted interaction score of 0.041. Both pairs are correctly classified using a decision threshold of 0.5.

The topology-only model is trained on the training split and used to score every pair in the corresponding held-out test split. The displayed neighborhoods are extracted from the same fold-specific drug–target–protein graph used to compute the topological descriptors for prediction. The negative example corresponds to the lowest-scoring pair in the test set, whereas the positive example is selected from the five highest-scoring test pairs. For the positive example, as for all positive test pairs, the corresponding DDI edge had been removed from the graph before topological feature computation, ensuring that the prediction is based solely on drug–target and protein–protein relations.

Figure 6a shows the documented positive DDI, whose local PPI context contains several proteins shared between the two drug neighborhoods. The drugs do not need to share the same direct targets for their neighborhoods to overlap: their respective targets may be connected through short PPI paths. Such connectivity provides a possible network-level basis for the interaction score assigned by the topology-only model.

**Fig. 6:**
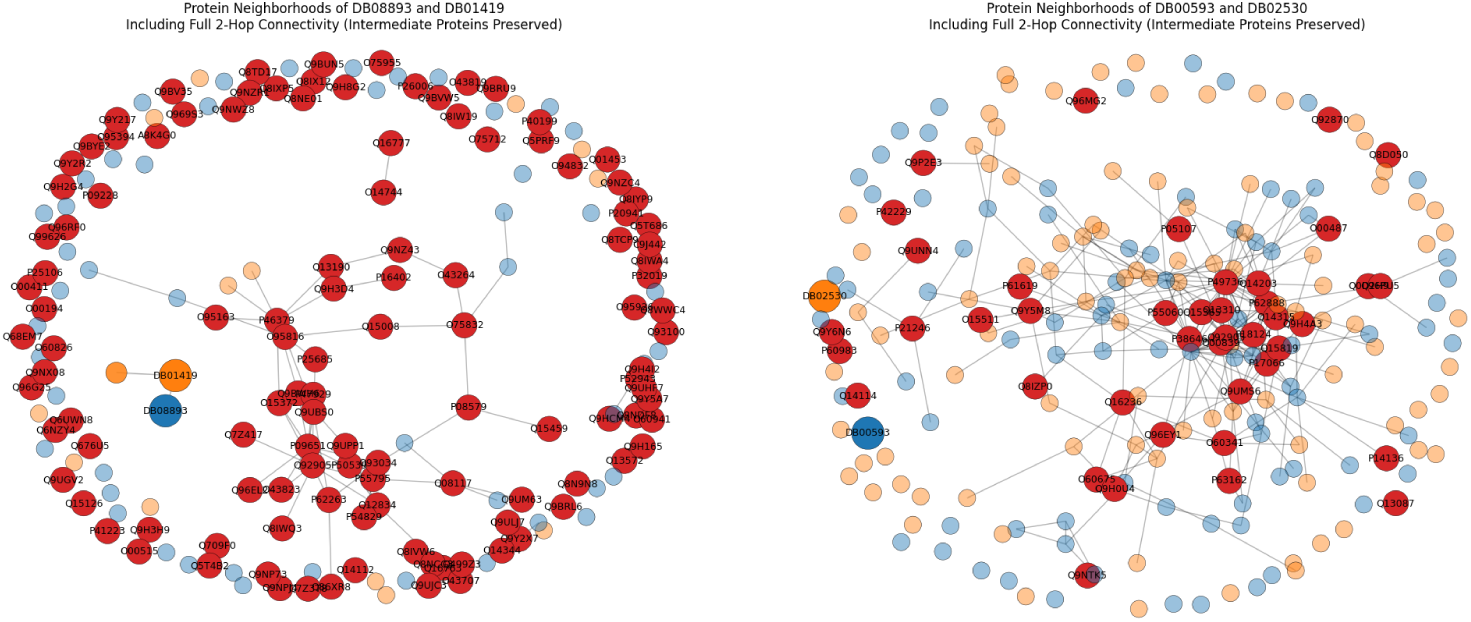
Qualitative examples of topology-only predictions. In each panel the two drugs are the largest nodes—Drug A in blue and Drug B in orange. Each drug’s direct protein targets are drawn in its own hue at high opacity and the surrounding proteins (within two hops of those targets) at lower opacity, with node size decreasing accordingly; proteins occurring in both drugs’ neighborhoods are highlighted in red. Every displayed node is a drug or one of these neighborhood proteins. Edges are all interactions among the displayed nodes in the induced subgraph.

In contrast, Figure 6b shows the curated reliable-negative pair, for which the two local neighborhoods contain fewer shared proteins and less overlap in their PPI context.

These examples illustrate the types of local network configurations captured by the topological descriptors. They provide qualitative intuition consistent with the multimodal-fusion results in Section 4.1, where topological information complements the MeSH-derived semantic representation.

## 6 Discussion

Our results highlight a practical challenge in computational DDI screening: the biomedical information available for individual drugs is uneven. Aggregate performance can remain high while masking weaker performance for sparsely annotated or unseen drugs. The depth-stratified and inductive analyses make this variation explicit.

### 1. Hierarchical supervision imbalance

Performance is generally lower when one or both drugs have shallow MeSH annotations, particularly in the inductive setting, where interactions involving unseen drugs must be predicted. Expanding the MeSH ontology scope from *Shallow* to *Intermediate* improves performance across nearly all interaction categories. In contrast, extending the scope from *Intermediate* to *Full* provides only limited and inconsistent additional benefit. Low–Low pairs remain among the most challenging categories. Within the present experiments, the *Intermediate* ontology scope therefore offers the most stable balance, retaining useful semantic information without depending on highly specific terms that are unavailable to drugs with shallower annotations.

### 2. When topology helps

In the transductive setting, adding DTP-KG topology through latent-space gating produces positive mean improvements over the MeSH-only baseline across all interaction categories. Under induction, the clearest gains occur for Low–Low and Low–Mid pairs, whereas improvements are smaller and less consistent when both drugs have Mid- or Deep-level annotations. This pattern suggests that graph-derived information is most useful when the semantic representation is limited by shallow ontology annotations.

The DTP-KG contains known training DDIs in addition to drug–target and protein–protein relations. The observed gains should therefore be interpreted as evidence for the contribution of graph-derived context as a whole, rather than being attributed exclusively to pathway-level or mechanistic information.

### 3. What the ablations reveal

Direct concatenation and global weighting produce only small, non-significant improvements over the MeSH-only baseline. Latent-space gating achieves the highest AUROC and is the only evaluated fusion strategy whose comparison with the MeSH-only baseline reaches the nominal threshold of *p <* 0.05. This result supports the use of instance-wise, dimension-specific fusion in the present setting, although the reported tests do not establish that latent gating is statistically superior to every alternative fusion strategy.

The grouped topology ablation shows that removing degree-based features produces the largest observed decrease in AUROC and F1. Removing clustering or centrality features leads to smaller decreases, indicating that these groups provide additional but comparatively smaller contributions to the topology-only model.

### 4. Interpretability and inductive behavior

The topology-only case studies illustrate local network configurations associated with high and low interaction scores, including differences in shared proteins and short PPI connections between drug neighborhoods. These selected examples provide qualitative intuition about the information captured by the topological descriptors, but they do not establish a causal explanation for individual DDIs or show that neighborhood overlap alone determines interaction status.

Performance decreases and variability increases in the inductive setting, consistent with the greater difficulty of predicting interactions involving unseen drugs. Fusion remains most beneficial in categories involving shallowly annotated drugs. The Deep–Deep category shows a negative mean AUROC difference under induction, but this difference is not statistically significant. The present experiments do not determine whether this pattern results from semantic distribution shift, graph structure, sample composition, or ordinary sampling variability.

#### Limitations and Future Work

- The DDI task is positive–unlabeled, and the present study relies on a curated set of reliable-negative pairs rather than a formal PU-learning objective. Positive-class downsampling avoids introducing additional unlabeled pairs as negatives, but it also exposes the model to only a subset of the available positive interactions during training. This may partly contribute to the limited improvement observed in Recall, although the effect was not evaluated directly. Future work should compare this strategy with class-weighted, cost-sensitive, and PU-aware objectives.
- The selected MeSH and graph modalities provide a common, structure-agnostic representation for small-molecule and biologic therapeutics. However, predictive performance is not reported separately for small molecule–small molecule, small molecule–biologic, and biologic–biologic pairs. The current results therefore establish shared representation coverage, but not therapeutic-class-specific effectiveness.
- The current approach assumes that each drug has sufficient ontology annotation and graph connectivity to construct the selected representations. It does not address a true cold-start setting in which a drug has neither usable semantic annotations nor known target or interaction information.
- We focus on lightweight, leakage-aware topological descriptors because they are computationally efficient and interpretable. Future work could compare this approach with end-to-end learning over the typed DTP-KG, including heterogeneous graph neural networks, and investigate additional modalities such as side-effect profiles, textual evidence, pathway annotations, or expression signatures.

## 7 Conclusions

We studied DDI prediction under heterogeneous biomedical knowledge availability and found that variation in MeSH annotation depth is associated with systematic differences in predictive performance that aggregate random-split evaluations can obscure. Expanding the ontology scope from *Shallow* to *Intermediate* provides substantial gains, whereas including Deep-level terms offers limited and inconsistent additional benefit.

Combining MeSH-derived semantics with graph-derived context from a Drug–Target–Protein knowledge graph through latent-space gating improves mean performance across all interaction categories in the transductive setting. Under induction, the clearest improvements occur for Low–Low and Low–Mid pairs, indicating that topological information is particularly useful when ontology annotations are shallow.

The resulting framework provides a shared, structure-agnostic representation that is compatible with both small-molecule and biologic therapeutics without requiring SMILES, molecular fingerprints, or amino-acid-sequence encodings. More broadly, the proposed depth-stratified evaluation offers a transparent way to assess DDI prediction under uneven semantic supervision and to identify conditions under which aggregate performance may conceal weaker generalization. The framework is intended to support scalable first-pass DDI prioritization for subsequent expert assessment.

## Declarations

## Consent for publication

Not applicable

## Availability of data and materials

The datasets analysed in this study were derived from the following publicly accessible resources:

- **DrugBank** (documented drug–drug interactions, drug–target interactions, and drug-level MeSH category assignments), DrugBank Online, release 5.1.13 (2 January 2025), https://go.drugbank.com/releases/5-1-13. DrugBank is distributed under licence and is therefore not redistributed in our repository; researchers with a valid DrugBank academic licence can reconstruct the analysis dataset from this release using our preprocessing scripts.
- **Medical Subject Headings (MeSH)**, U.S. National Library of Medicine, 2025 descriptor release (desc2025.xml), https://nlmpubs.nlm.nih.gov/projects/mesh/MESH_FILES/xmlmesh/ (download portal: https://www.nlm.nih.gov/databases/download/mesh.html). MeSH is in the public domain.
- **UniProt** (protein–protein interactions and annotations), UniProt Knowledgebase, release 2025 03, https://ftp.uniprot.org/pub/databases/uniprot/previous_releases/release-2025_03/. Proteins are identified by UniProt accession numbers; UniProt data are available under the Creative Commons Attribution 4.0 licence.
- **Reliable negative DDI pairs**, obtained from Additional file 2 (Table S3) of Zheng et al. [5] (DDI-PULearn), BMC Bioinformatics, https://doi.org/10.1186/s12859-019-3214-6.

The source code, configuration files, and analysis and preprocessing scripts required to reproduce all results are openly available at https://github.com/Golnazthr/DTP-KG.

## Ethics approval

Not applicable

## Authors’ contributions

AY contributed to the methodology, software implementation, formal analysis, validation, visualization, and writing of the original draft. SS contributed to the methodology. GT contributed to the conceptualization, supervision and funding acquisition. AY and GT reviewed and edited the manuscript. All authors read and approved the final manuscript.

